# Microglia and Chek2 contribute to sex-specific organization of the adult zebrafish brain

**DOI:** 10.1101/2025.08.15.670359

**Authors:** Paloma Bravo, Florence L. Marlow

## Abstract

Sex specific differences in size and distribution of cell types have been observed in mammalian brains. How sex-specific differences in the brain are established and to what extent sexual dimorphism contributes to sex-biased neurodevelopment and neurological disorders is not well understood. Microglia are the resident immune cells of the nervous system and have been implicated in masculinizing the mammalian brain and refining neural connections to promote remodeling of neural circuitry, yet their contributions to developmental brain patterning and plasticity in zebrafish remains unclear. Here, we report anatomical and cellular differences between juvenile brains and adult female and male brains. Leveraging the plasticity of the zebrafish female brain and genetic models lacking microglia and tumor suppressor factors, we provide insight into the mechanisms that establish sex-specific brain dimorphism in zebrafish. Specifically, we identified sexually dimorphic features in the adult zebrafish brain that depend on microglia and Chek2, which may have broader implications and represent therapeutic targets for sex-biased neurological disorders.

**Plain language summary:** Males and females of species can have significant differences in appearance, including differences in size, color, or sex specific anatomical structures. In addition to overt morphological differences, sex specific differences in size and distribution of cell types have been observed in mammalian brains. How these sex-specific differences in the brain are established and to what extent these differences contribute to sex-specific neurodevelopment and neurological disorders that differentially impact males and females is not well understood. Despite an incomplete picture of the mechanisms regulating sex-specific development, some of the cell types involved include microglia. Microglia are the resident immune cells of the nervous system and have been implicated in promoting features that are typical in the male mammalian brain. Specifically, microglia may refine neural connections and promote remodeling of neural circuitry and influence sex-specific behaviors. The contributions of microglia to developmental brain patterning and plasticity in zebrafish remain unclear. Here, we report anatomical and cellular differences between juvenile brains and adult female and male brains. Leveraging zebrafish genetic models lacking microglia and tumor suppressor factors, and the unique plasticity of the zebrafish female brain, we investigated and provide insight into the mechanisms that establish sex-specific brain differences in zebrafish. Specifically, we identified sexually distinct features in the adult zebrafish brain that depend on microglia and the tumor suppressor Chek2. If these or similar mechanisms operate in other species, our findings may have broader implications for sex-specific brain development and represent therapeutic targets for sex-biased neurological disorders.

**Highlights:**

- Tissue clearing and immunostaining of juvenile and adult whole-mount zebrafish brains allows analysis of sex differences.
- Anatomical and cellular sexual dimorphism in the adult vertebrate brain appears after gonadal sex differentiation.
- Sexual dimorphism in the adult brain is driven by differences in cell death regulation.
- Microglia colonization of brain areas involved in courtship is sexually dimorphic.
- Microglia involvement in establishing sex-specific differences in the adult brain.

## Background

Females and males within species share almost all genetic information and yet can develop considerably diverse morphologies at the whole animal and tissue level. In some cases, with distinct functions and morphologies that cause sex-specific susceptibility to diseases (Lopes-Ramos et al., 2020; Oliva et al., 2020; Yang et al., 2006). Anatomical, cellular, and molecular differences have been observed between female-typical and male-typical brains of most vertebrates, including humans (Agate et al., 2003; Dewing et al., 2003; Gillies et al., 2014; Kipp et al., 2012; Li and Singh, 2014; Ramien et al., 2016; Scholz et al., 2006). These differences are associated with sex-specific gene expression profiles, and variation in the levels of sexual hormones, including androgens and estrogens, and are thought to be established and maintained by sex hormone responsive cells that reside within each tissue (Agate et al., 2003; Carruth et al., 2002; Dewing et al., 2003; Maekawa et al., 2014; Peterson et al., 2013; Scholz et al., 2006).

In sexually reproducing animals, an initially sexually indeterminant individual acquires sex specific reproductive organs and secondary sex traits during sex determination and differentiation. The mechanisms of sex determination vary between species and can be regulated by dedicated sex chromosomes, as in mammals, or multigenic determination systems, as in zebrafish, and environmental factors (Capel, 2017; Nagahama et al., 2021; Wilhelm et al., 2025). In most species, the earliest stages of gonad development and throughout reproductive life, intragonadal signals between germline and somatic gonad and extragonadal signaling ensures proper reproductive development and function. This includes production of the hormones necessary for proper sexual differentiation of the gonads and gamete production, and development of the secondary sex traits that are required for fertility (Estermann et al., 2020a; Estermann et al., 2020b; Nagahama et al., 2021).

Disruption of these signals can cause misalignment of germ and somatic cell sex within the gonad and throughout the body (Donohoue, 2020). For example, in mammals, exposure to gonadal hormones at critical stages of development patterns the sex-typical circuits, including neuronal numbers, cell types and connectivity, that determine social behaviors in adults (MacLusky and Naftolin, 1981; McCarthy, 2008; Simerly, 2002). In mice, estradiol is the key factor driving sexual differentiation of the neonatal brain. In males, a spike of testosterone produced after birth provides a source of estradiol that has both cell autonomous and nonautonomous influences on sex specific gene expression and neuronal subtypes and numbers (Gegenhuber et al., 2022). In female mice, estradiol maintains active repression of genes that promote male-typical patterns (Nugent et al., 2015; Oberlander and Woolley, 2016; Tabatadze et al., 2015). After development, and throughout adulthood, sex hormones continue to play important roles in maintaining reproductive tissues and behaviors (Chakravarthi et al., 2021; Downs and Wise, 2009; Marshall et al., 1991). In animal models and in humans both reproductive and neurological disorders can arise from disruption of sexual development (Joel et al., 2016; López-Ojeda and Hurley, 2021; McCarthy, 2016). Thus, it is important to understand how gonad differentiation and function are coordinated with development of the brain, especially establishment and maintenance of the sex-specific brain features that can influence function and behaviors throughout life.

Although the mechanisms that establish sexually dimorphic patterns and circuits are not fully understood, microglia, the immune cells of the brain, have been implicated in regulating neurogenesis, oligodendrogenesis, cell proliferation, and pruning synapsis but the specific contributions of microglia to sculpting neural circuits during development are still unclear (Casano et al., 2016; Lenz and Nelson, 2018; Li et al., 2019; Lopez-Atalaya et al., 2018; Nelson et al., 2019; O’Keeffe et al., 2025). Thus, microglia have been proposed to play a role in shaping the brain during development and/or adulthood. Accordingly, studies have identified sexual dimorphism in microglia numbers and regional distribution between the brains of male and female mice (Lenz and McCarthy, 2015). Notably, in addition to producing prostaglandins (Nugent et al., 2009; Wright and McCarthy, 2009; Zhang et al., 2009), lipid compounds that function as hormones in regulating bodily functions, and expressing prostaglandin receptors (Niraula et al., 2023), microglia are also highly responsive to sex hormones, including estrogens (Baker et al., 2004; Loiola et al., 2019; Morale et al., 2006; Perez-Pouchoulen et al., 2019; Xu et al., 2016a). Accordingly, prominent sex-specific differences in microglia have been documented in early development of mammals, including differential colonization of the developing brain based on estrogen or androgen exposure and region-specific microglia activation states that extends throughout lifespan (Nelson et al., 2019). Furthermore, microglia and prostaglandins contribute to masculinization of the mouse brain - specifically to development of the dendritic spine density associated with male specific sexual behaviors (Lenz et al., 2013a). Pharmacological inhibition of PGE2 results in diminished microglia numbers and cellular morphologies that resemble those observed in females (Lenz et al., 2013b). Conversely, stimulation with PGE2 results in females with microglia numbers and cellular morphologies like those of males and requires microglia activation.

Like mice and other sexually reproducing animals, zebrafish males and females are identifiable by distinct sex-specific morphological features, mating behaviors, and sex-specific neural pathways associated with those behaviors (Kimchi et al., 2007; Yabuki et al., 2016). For example, male courtship behaviors are stimulated by prostaglandin F2 alpha (PGF2a) produced by females (Yabuki et al., 2016). Although the mechanisms underlying formation of these sexually dimorphic neural pathways is not fully understood, the ability of zebrafish to completely switch their phenotypic sex and behaviors from female to male in response to environmental or genetic factors that impair fertility suggests that the female brain retains developmental plasticity (Beer and Draper, 2013; Dranow et al., 2016; Dranow et al., 2013; Draper et al., 2007; Rodriguez-Mari and Postlethwait, 2011; Shive et al., 2010). Females that switch sex adopt male-specific courtship behaviors whether sex reversal occurs developmentally, soon after gonadal differentiation or in adulthood. Thus, female to male sex reversal is a unique system to study plasticity of the developing and mature brain. In some fish species, transitioning of sex-specific behavioral traits precedes detectable changes in circulating hormones (Dodd et al., 2019). This observation led to a “sequential hermaphroditism model” in which sex change is initiated by the brain in response to interpretation of cues from the social environment. In contrast, genetic data in zebrafish indicate that sex reversal is initiated in the ovary. For example, disruption of *nanos3,* a conserved RNA binding protein that is specifically expressed in the germline where it regulates GSC renewal and maintenance, causes sex-reversal in adulthood after depletion of the nonrenewing pool of oocytes (Beer and Draper, 2013; Draper et al., 2007). Moreover, female fish with mutations in the conserved, oocyte-expressed ligand, *bmp15,* never produce mature eggs and undergo ovary to testis reversal, in part due to deficits in estrogen production (Dranow et al., 2013; Zhai et al., 2023), and through mechanisms involving activation of macrophages by cytokines expressed by somatic gonadal cells (Bravo et al., 2023). However, whether sexual dimorphism in microglia colonization and morphology similar to that observed in mammals also occurs within the developing and adult zebrafish brain and potential microglia contributions to brain plasticity remain to be determined.

In this work, we report anatomical and cellular features of developing and adult female and male zebrafish brains. We leveraged the plasticity of the zebrafish female brain and genetic models lacking microglia and tumor suppressor factors to investigate the mechanisms that establish sex-specific brain dimorphism in zebrafish. Specifically, we identified sexually dimorphic features in the adult zebrafish brain that depend on microglia and Chek2, which may have broader implications and represent therapeutic targets for sex-biased neurological disorders.

## Methods

### Animals

Zebrafish mutant lines used for experiments performed within this work were: *bmp15^uc31^, chek2^sa20350^, irf8^st96^, nanos3^fh49^, and tp53^zdf1^* (Beer and Draper, 2013; Berghmans et al., 2005; Dranow et al., 2016; Kettleborough et al., 2013; Shiau et al., 2015a). Experimental fish referred to as ‘adults’ in whole-mount immunohistochemistry and live tissue images were 3 months old or older. Juvenile fish used for brain analysis were either 28dpf or 35dpf. Standard husbandry and care conditions were used for maintenance of all zebrafish. All protocols and procedures were performed following guidelines from the National Institutes of Health and approved by the Icahn School of Medicine at Mount Sinai Institutional (ISMMS) Animal Care and Use Committees (IACUC, 2017-0114).

### Genotyping

Gonad tissue, fin clip or trunks were lysed in an alkaline lysis buffer (25 mM NaOH and 0.2 mM EDTA pH 12) to extract genomic DNA (gDNA), then heated for 20 min at 95° and cooled at 4°C before adding neutralization buffer (20 mM Tris-HCl and 0.1 mM EDTA pH 8.1) (Truett et al., 2000). gDNA was used as template for PCR amplification, digested if needed with a restriction enzyme, and resolved in a 3% 1:1 MetaPhor/agarose gel for genomic diagnosis. Genotypes were confirmed by sequencing.

### Microscopy imaging of brain and gonadal tissues

Live anatomical tissue images were acquired with a Stemi stereo microscope (ZEISS Microscopy, Germany). Dissected gonads and brains were imaged using a Zoom.V16 microscope (ZEISS Microscopy, Germany).

Brain images acquired after immunofluorescence staining were obtained using image tiling on a light-sheet UltraMicroscope II (Miltenyi Biotec, Germany) equipped with a Neo 5.5 sCMOS camera (Andor, Ireland). Samples were imaged with Olympus magnification lenses of 4X or 12.6X (1X zoom), using the three light sheet configuration from a single side, with the horizontal focus centered in the middle of the field of view, and a light sheet width of 60–100% (adjusted depending on sample size to ensure even illumination in the y-axis). Spacing of Z slices was 3 or 5 µm. Samples were illuminated with 568nm and 640nm lasers (Coherent, Germany). The chromatic correction module on the instrument was used to ensure both channels were in focus.

### Neutral red staining of zebrafish embryos

Embryos were stained with neutral red solution to mark microglia as previously described (Herbomel et al., 2001). Briefly, embryos were kept alive in a solution of embryo media with 0.003% propylthiouracil (PTU-water) that was replaced every 1-2 days to prevent the appearance of pigmentation. At the experimental age, embryos were incubated in the dark for 5h at 28.5°C in a solution of 40ml PTU-water with 20ul of neutral red solution (Sigma Aldrich). Embryos were then washed at least 2 times with embryos media and kept at 28.5°C until imaging.

### Juvenile and adult brain dissections and fixation

Fish were anesthetized with a lethal dose of tricaine (MS-22) (400mg/l). Brains were dissected prior to or after fixation. Briefly, the head was removed with a razorblade and placed upside down, exposing the ventral side of the head. With forceps, the jaw and soft tissue within the head were removed until the bones of the skull were visible. Using spring microdissection scissors both optic nerves connecting the eyes to the optic chiasm were severed and the rest of the face bones and eyes were pulled away. The bones of the skull were removed, starting in the most anterior region of the head (olfactory bulb) and advancing to the posterior part of the head until the whole brain was exposed, and most brain anatomical structures were visible. Dissected brains were placed in 4% PFA overnight, then washed with PBS, dehydrated in MeOH, and kept at -20°C until use.

### Whole-mount immunofluorescence, tissue embedding, and clearing

After fixation, samples were stored in MeOH at -20C overnight or until use. Whole tissue staining, embedding, and clearing was performed as described in detail in (Lindsey et al., 2018). All immunofluorescence steps were done in the dark, at 4°C, in a 12-well plate where individual samples were placed in mesh baskets and transferred from one well to the next as follows: 1) rehydration with 0.3% PBSTx (0.3% Triton X-100 in PBS), 2) permeabilization with 5% DMSO in 1% PBSTx, 3) incubation overnight in blocking buffer (2% normal goat serum;1% bovine serum albumin in 0.3% PBSTx), 4) primary antibody incubation in blocking buffer for 7 days, 5) wash with 0.3% PBSTx, 8) secondary antibody incubation in 0.3% PBSTx for 7 days, 9) nucleic acid stain (Sytox 1:2000 in 0.3% PBSTx) overnight, and 10) wash in 0.3% PBSTx. Following staining, brains were washed in milliQ water and embedded into 1% low-melting agarose as described in(Lindsey et al., 2018). Briefly, each brain was placed in the center of a well in a 6-well plate filled with filtered 1% low-melting agarose and let cool down at 4° C for at least an hour before trimming excess. Clearing of the embedded tissues was performed at room temperature in a fume hood and consisted of several consecutive incubations in 100% MeOH, followed by incubations in BABB (1:2 benzyl alcohol/benzyl benzoate) (Sigma Aldrich). Images of cleared, embedded tissues were acquired using a LaVision Ultramicroscope II light sheet microscope and analyzed/processed using Imaris Viewer (Bitplane).

### Brain region analyses

Anatomical measurements of the specified regions of the brain were made in Imaris Viewer (BitPlane) using the section view with normal and extended crosshair. Analysis of microglia densities was done by manually counting Sytox-positive cells and 4c4-positive cells in 200×200 pixel size squares of 3 difference areas in a 3×3 grid placed over a 5-slices MAX projection.

### Statistical Analysis

Statistical analyses of adult brain anatomical comparisons were performed using ordinary TWO-way ANOVA with multiple comparisons (uncorrected Fisher LSD) and 95% CI. P-values: ns ≥ 0.05, *P ≤ 0.05, **P ≤ 0.01, ***P ≤ 0.001. Statistical analyses of microglia cell density comparisons were done by unpaired t-test with Welch’s correction and 95% CI. P-values: ns ≥ 0.05, *P ≤ 0.05, **P ≤ 0.01, ***P ≤ 0.001. Analysis and statistical graphical representations were done in Prism (GraphPad Software, Inc.).

## Results

### Morphological differences in the juvenile and adult zebrafish brain

In sexually reproducing species, mating involves sex-specific behaviors and circuits that mediate gamete release and sex-specific reproductive behaviors. Where examined, differences in brain anatomy have been documented in the brain regions that mediate reproductive behaviors. Accordingly, courtship in zebrafish involves defined sex-specific adult behaviors that are mediated by the brain regions and nuclei that influence gamete release and mating (Kermen et al., 2013; Yabuki et al., 2016). We and others have shown that zebrafish that undergo female-to-male sex reversal adopt male mating behaviors, but whether zebrafish exhibit sexual dimorphism in brain anatomy has not been reported (Bravo et al., 2023; Dranow et al., 2016; Zhai et al., 2023). Therefore, we investigated whether morphological or cellular differences between female and male zebrafish brains are detectable in brain regions that regulate courtship and mating behaviors (Fig. 1A-1D). Specifically, we performed whole mount immunohistochemistry, light sheet microscopy, and measured the brain regions involved in mating, the telencephalon (Te), optic tectum (OpT), and hypothalamus (HT)(Kermen et al., 2013; Yabuki et al., 2016). We examined juveniles (28dpf and 35dpf), and adult (>3months) male and female zebrafish brains (Fig. 1E), to determine if these regions were sexually dimorphic in size. Because sexual dimorphism of the brain is influenced by sex hormones in other species, and because lab strains develop a sexually indeterminant gonad before developing a juvenile ovary, we expected juvenile brains would not be dimorphic. As anticipated for stages prior to sex determination, no dimorphism in volume of the Te, OpT, or HT of juvenile brains were apparent (Fig. 1F).

**Figure 1.**
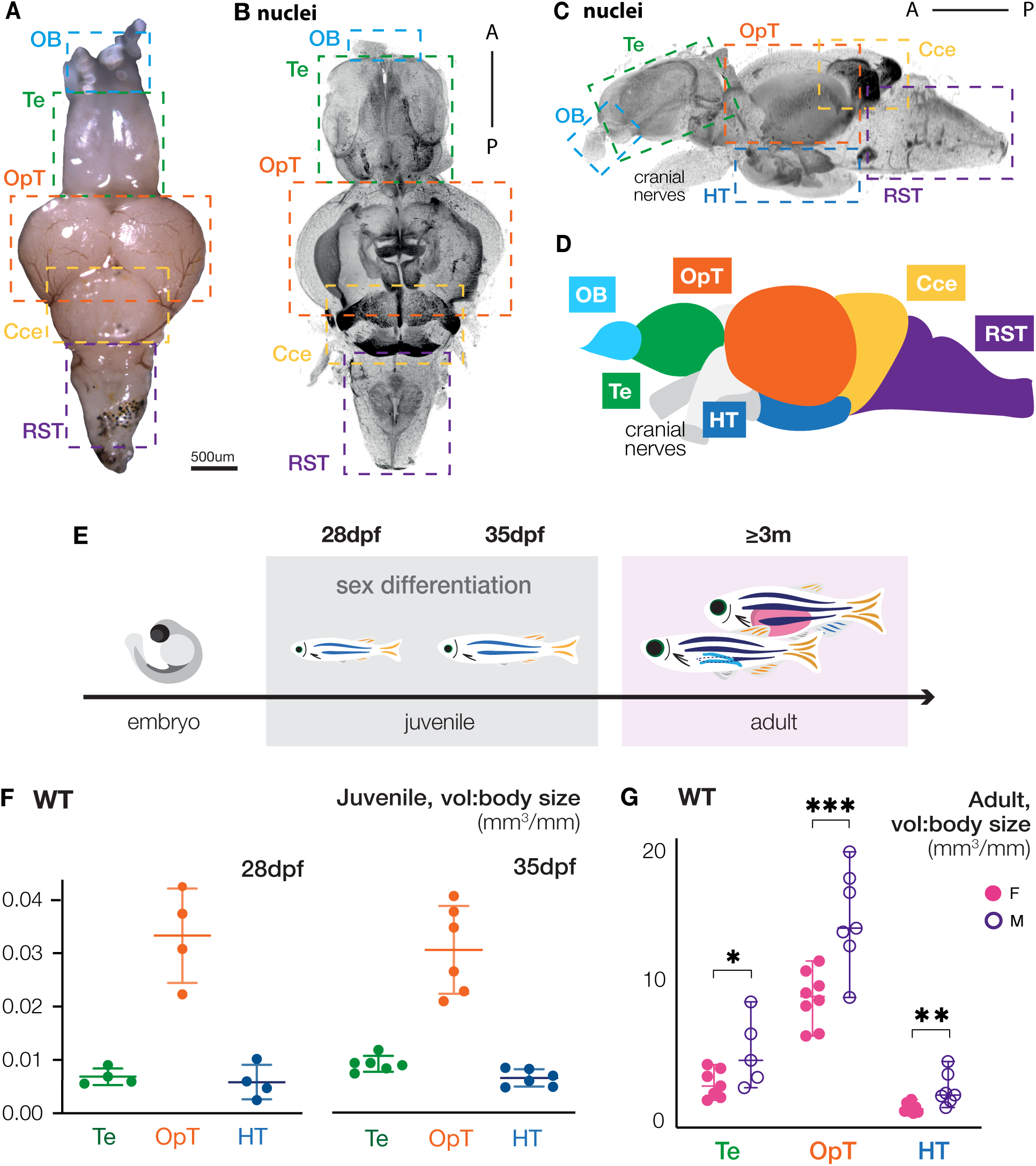
Morphological differences between female and male adult brains emerge after sex determination. **(A-D)** Adult **(A)** live tissue, **(B and C)** immunostained image, and **(D)** illustration of the zebrafish brain with color-coded, dotted-line boxes indicating anatomical structures. **(E)** Zebrafish developmental timeline with experimental stages used for brain analyses. **(F and G)** WT brain volume/body quantification of indicated regions of **(F)** adult females and males, and **(G)** indeterminant (28dpf) and juvenile gonads (35dpf). OB: olfactory bulb; Te: telencephalon; OpT: optic tectum; Cce: cerebellum; RST: reticulospinal tract; HT: hypothalamus. Statistical analysis: ordinary Two-way ANOVA with 95% CI, and Fisher LSD test for multiple comparisons. P-values: ns≥ 0.05, *<0.05, **<0.01, ***<0.001.

Next, we examined the brains of fully differentiated, sexually mature adult males and females. In contrast to brains of sexually immature juveniles, the brain regions associated with sex-specific courtship behaviors – the Te, OpT, and HT – were sexually dimorphic and were larger in males than in females (Fig. 1G), indicating that sexual dimorphism in the brain arises only after gonadal sex is fully developed. This finding aligns with previous research describing anatomical, physiological, and behavioral effects on mammalian brains exposed to sex hormones during development (Gegenhuber et al., 2022; Juntti et al., 2010; Wu et al., 2009).

We hypothesized that the larger size of the Te, OpT, and HT in male brains could be a consequence of prolonged growth of the male brain relative to the female brain, or alternatively, due to equivalent growth but different rates of cell death between the sexes. Programmed cell death through apoptosis has been extensively studied as a mechanism to refine tissue morphologies during development (Ameisen and Ameisen, 2002; Jacobson et al., 1997; Kerr et al., 1972). In the developing nervous system, half of the neurons that are born ultimately die and are eliminated through clearance processes involving immune cells Dekkers et al., 2013; Pfisterer et al., 2017). To determine if the differences in volumes between the female and male brain regions were due to increased proliferation in males or increased cell death in females, we analyzed brains of both sexes in the absence of Chek2. Chek2 is a tumor suppressor factor with essential roles in the activation of Tp53- and Tp63-dependent cell death pathways, and Chek2 ablation has been previously shown to suppress cell loss and tissue damage phenotypes (Bolcun-Filas et al., 2014; Bravo et al., 2023; Chihiro et al., 2023). As expected, based on their normal mating behaviors, the Te, HT, and OpT of heterozygous *chek2* fish, were sexually dimorphic like wild type (Fig. 2A-2C). However, while the Te and OpT were larger in males, the HT of *chek2* heterozygous females was larger than in males (Fig. 2C). In contrast to the sexual dimorphism observed in wild-type and *chek2* heterozygous adult brains, we observed no sex-specific differences in the Te, OpT, and HT of adult *chek2* mutant zebrafish brains (Fig. 2A-2C). Because no sexual dimorphism of the Te, OpT, and HT was detected among *chek2* mutants, we conclude that Chek2-mediated cell death rather than enhanced neurogenesis in males is the main driver of the smaller size of these brain regions in females.

**Figure 2.**
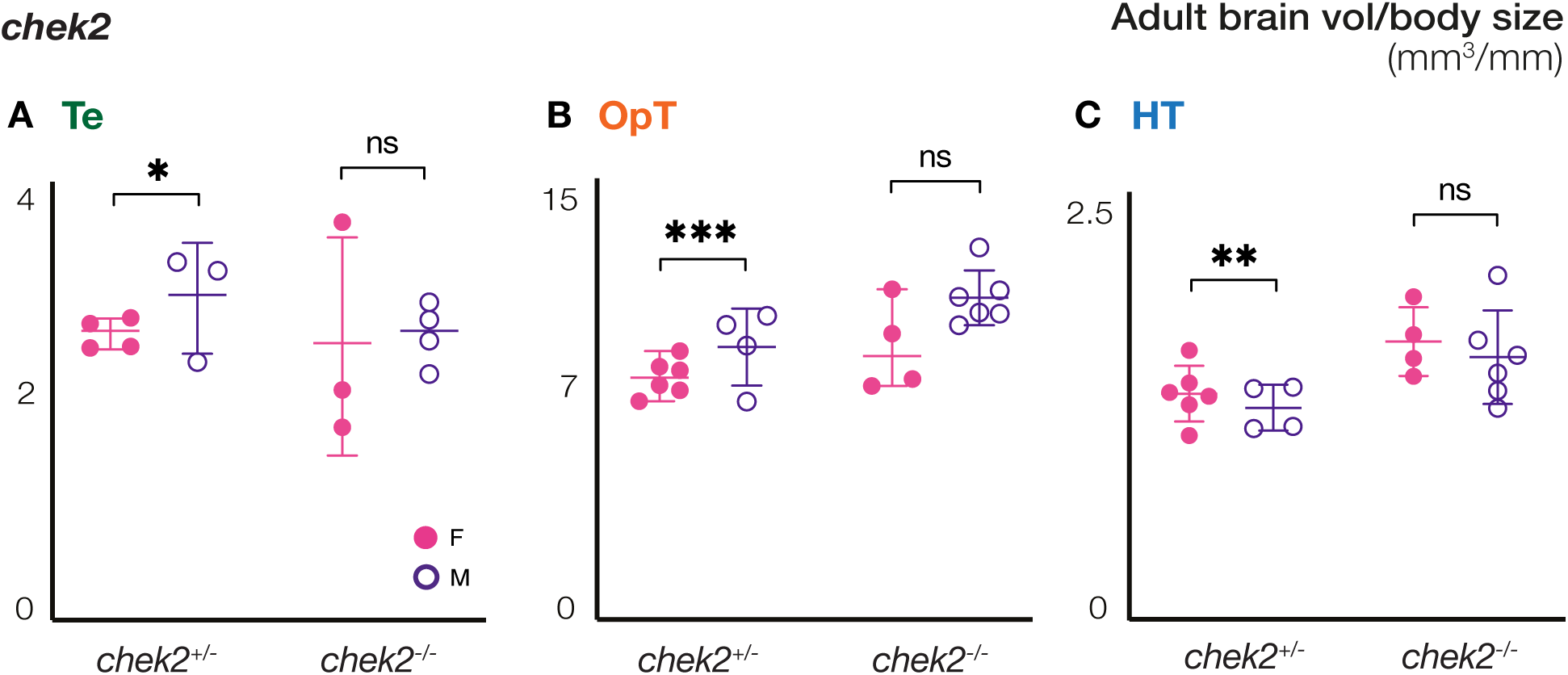
Chek2 contribution to regional size differences between female and male adult brains. **(A-C)** Comparison of volume/body sizes of the **(A)** Te, **(B)** OpT, and **(C)** HT of adult female and male brains of the indicated genotypes. Te: telencephalon; OpT: optic tectum; HT: hypothalamus. Statistical analysis: ordinary Two-way ANOVA with 95% CI, and Fisher LSD test for multiple comparisons. P-values: ns≥ 0.05, *<0.05, **<0.01, ***<0.001.

### Morphological and cellular brain organization after female-to-male sex-reversal

In zebrafish, female-to-male sex-reversal can be induced by a variety of environmental or genetic factors (Aharon and Marlow, 2021; Dranow et al., 2013; Kossack and Draper, 2019). In the absence of Bmp15, a conserved regulator of follicle progression, the zebrafish ovary undergoes oocyte loss, masculinization, and is remodeled into a testis (Dranow et al., 2016) (Suppl. Fig. 1A). Sex reversal due to loss of Bmp15 takes place between day 35 to 85 post fertilization and is mediated by a mechanism that involves cell death and Csf1a-mediated activation of gonadal macrophages (Bravo et al., 2023). This masculinization extends beyond the gonad to all secondary sex-specific traits, including pigmentation, fin morphology, and behavior (Dranow et al., 2016). Presumably, acquisition of new sex-specific behavioral pathways, including mating, by sex-reversed *bmp15* mutants involves changes in the brain, but this possibility has not been examined. Therefore, we analyzed the size of the Te, OpT, and HT of adult brains of *bmp15* heterozygotes and compared them to brains of non-sibling adult wild type of similar age. As expected, and like wild-type, the Te, OpT, and HT were sexually dimorphic in size between female and male *bmp15* heterozygotes (Fig. 3A-3C). Interestingly, the volume-to-body size ratio was statistically different between WT and heterozygotes for the OpT: both heterozygous female and male OpT were larger than WT of similar age and size (Suppl. Fig. 1B). The regional volumes of *bmp15* homozygous mutant male brains were more comparable in size to those of heterozygous males than heterozygous females (Fig. 3A-3C). Because adult mutant brains were analyzed at stages after female-to-male sex reversal occurs, direct differentiating males and sex-reversed females could not be distinguished based on gonad histology or other anatomical features. Nonetheless, these results suggest that sex-reversed *bmp15* mutants acquire the morphology of direct developing males in areas of the brain associated with sex-specific behaviors.

**Figure 3.**
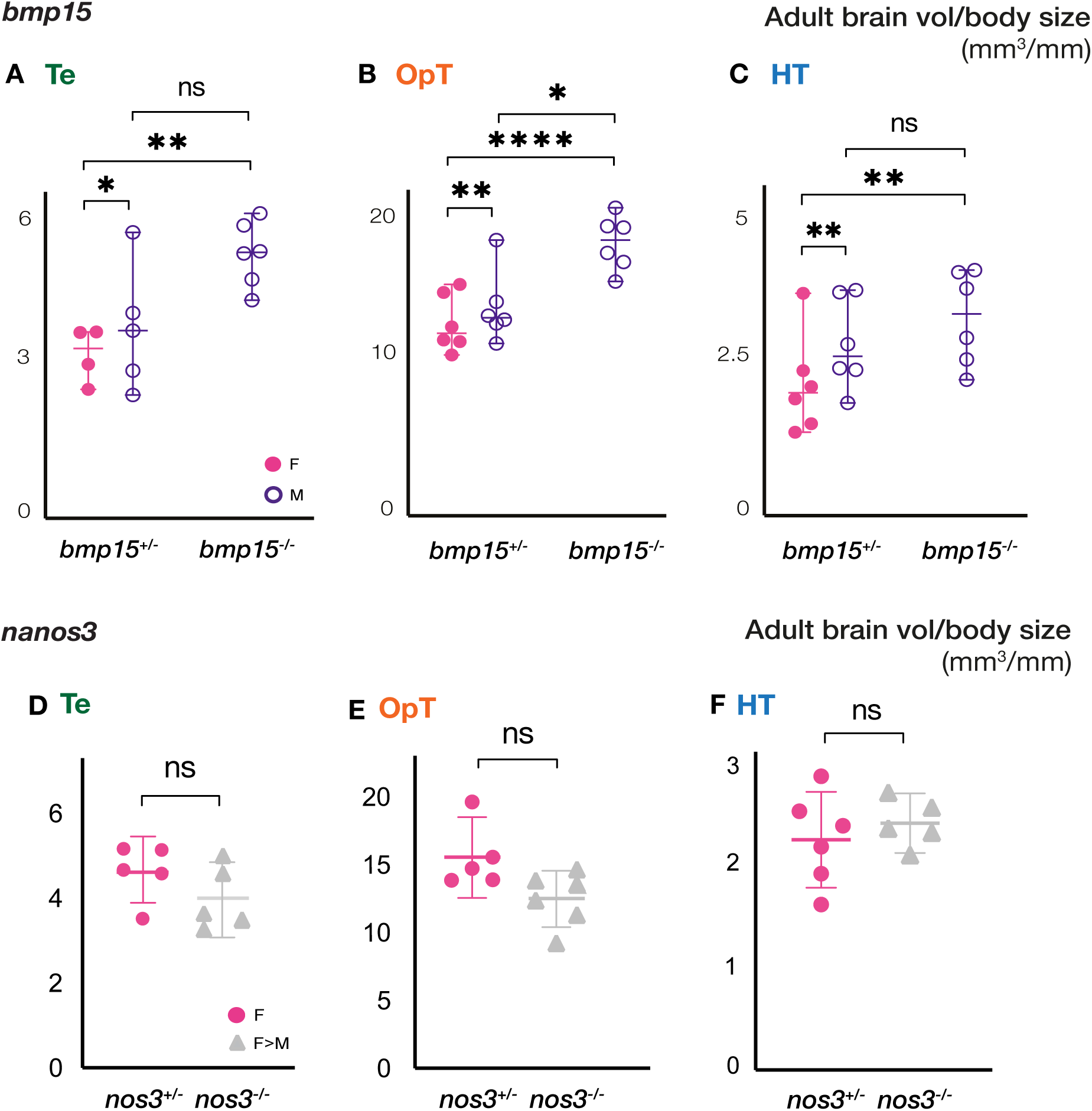
Adult neuroanatomy of *bmp15* but not *nanos3* mutants resembles that of direct-developing males. **(A-C)** Comparison of volumes of **(A)** Te, **(B)** OpT, and **(C)** HT of adult females and males of the indicated genotypes. **(D-F)** Comparison of volumes of **(D)** Te, **(E)** OpT, and **(F)** HT of adult females and female-male sex reversed mutants of the indicated genotypes. Te: telencephalon; OpT: optic tectum; HT: hypothalamus. Statistical analysis: **(A-C)** ordinary Two-way ANOVA with 95% CI, and Fisher LSD test for multiple comparisons, **(D-F)** unpaired t-test with Welch’s correction with 95% CI. P-values: ns≥ 0.05, *<0.05, **<0.01, ****<0.0001.

Like mutation of *bmp15,* loss of Nanos3 triggers female to male sex reversal; however, reversal occurs later, in adult females that have depleted their nonrenewing pool of oocytes (Beer and Draper, 2013). Because sex-reversal of *nanos3* mutants fish occurs after a mature ovary has developed, we were able to identify and track previously mated *nanos3* mutant females and compare their brains after sex reversal. In contrast to sex reversal in young fish, no anatomical differences in the brain areas involved in mating (Te, OpT, and HT) were detected between *nanos3^+/-^*females and sex-reversed *nanos3^-/-^* mutants (Fig. 3D-3F). Females of all genotypes were housed individually for the mating trials, and thus were significantly larger than, and could not be compared to, their male siblings (Suppl. Fig. 1C-1E). The comparable sizes between brain areas of *nanos3^+/-^* females and sex-reversed *nanos3^-/-^* mutants suggests that there are no gross anatomical changes in the brain when sex reversal occurs in mature adults; however, molecular or cellular changes in microglia or other populations may contribute to establishing male-specific behaviors.

### Microglia colonization of the larval and adult brain in the absence of Tp53

We previously showed that eliminating all macrophages, including microglia, blocks sex reversal of *bmp15* mutants (Bravo et al., 2023). In this context, blocking ovary to testis reversal prevents reversal of all sexually dimorphic body features and behaviors, suggesting reversal of the gonad precedes and is prerequisite for female to male sex reversal (Bravo et al., 2023). This contrasts with protandrous species that can reverse sex from male to female and in which sex specific behaviors and brain sex reversal precedes and can be uncoupled from gonad sex (Dodd et al., 2019; Parker et al., 2024). In zebrafish, global loss of *irf8* eliminates macrophages and microglia, thus distinct contributions of gonadal macrophages and microglia to sex reversal cannot be easily determined. Prior work showed that Tp53-dependent apoptosis is required for microglia colonization of the embryonic zebrafish brain (Casano et al., 2016; Xu et al., 2016b). Using morpholinos targeting Tp53 and a hindbrain specific nitroreductase (NTR) ablation system, cell death induced by NTR ablation or UV irradiation resulted in increased microglia colonization in a Tp53-dependent manner (Casano et al., 2016). Moreover, in this context, altered microglia numbers persisted weeks after ablation; however, whether these changes persist in adulthood was not reported. Thus, we attempted to dissect macrophage and microglia contributions to sex reversal by blocking macrophage colonization of tissues (Casano et al., 2016; Ferrero et al., 2020; Iyer et al., 2020; Kierdorf et al., 2013; Li et al., 2011; Lou et al., 2022; Shiau et al., 2015b; Wu et al., 2018; Xu et al., 2016b). To determine if blocking Tp53-mediated microglia colonization could be leveraged to eliminate microglia from the adult zebrafish brain and to explore microglia involvement during sex reversal, we examined microglia colonization in the brains *tp53* mutant larvae and adults. In contrast to the morpholino studies, microglia were observed in the brains of Tp53 maternal-zygotic mutant larvae (Fig. 4A-4B) and adults (Fig. 4C-4E). Therefore, determining microglia specific contributions to establishing sex-specific behaviors after female-to-male sex reversal awaits development of tools to specifically ablate microglia without disrupting macrophages.

**Figure 4.**
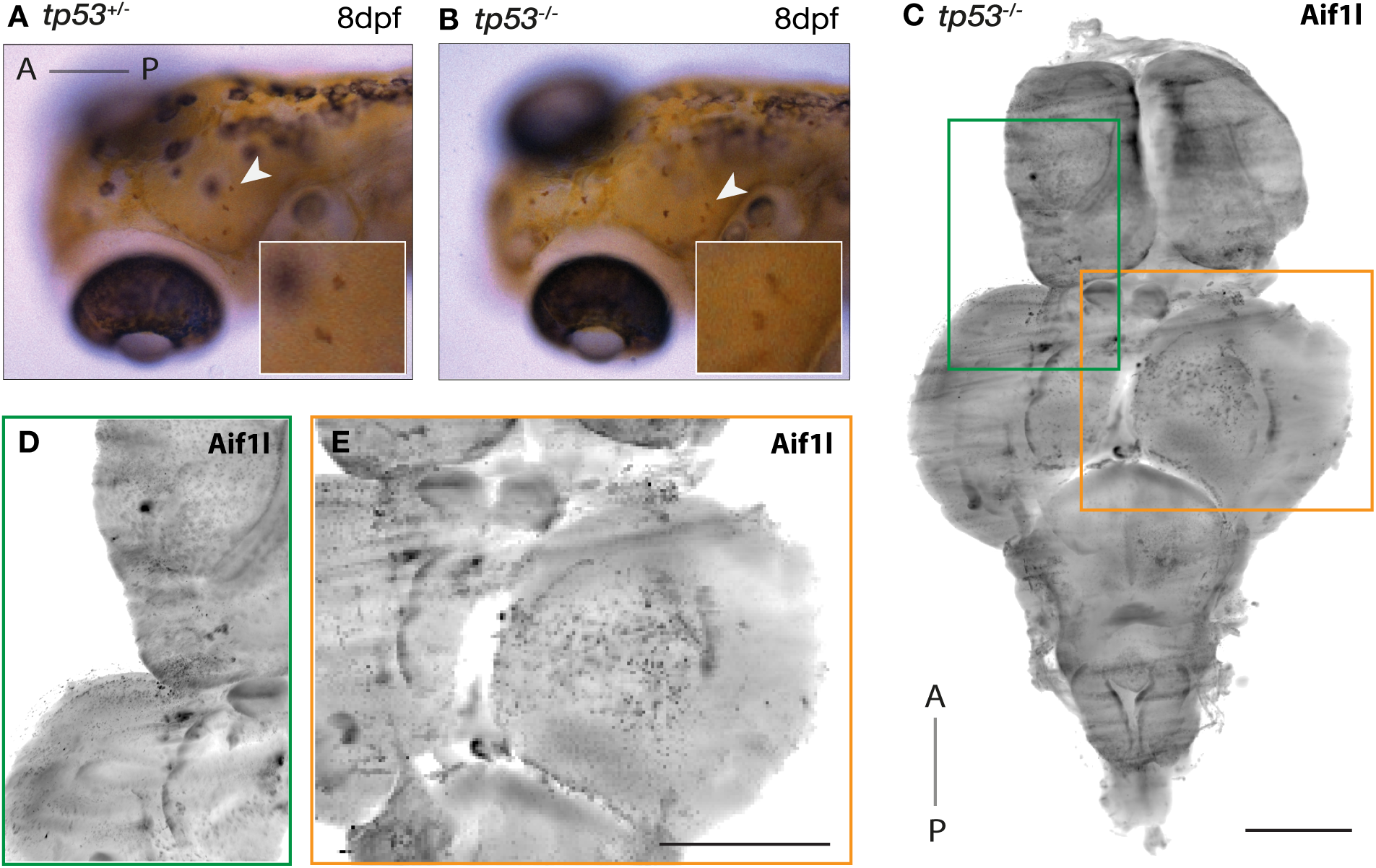
Microglia colonize the zebrafish brain in the absence of Tp53. **(A and B)** 8dpf **(A)** *tp53^+/-^* and **(B)** *tp53^-/-^*embryos stained with neutral red to label microglia. White arrow and boxes are magnified views showing neutral red-positive cells. **(C-E)** Adult *tp53* mutant brain stained for 4c4 (microglia). Scale bars: 1mm. Boxes correspond to magnified regions of the **(D)** telencephalon and **(E)** optic tectum.

### Microglia contributions to sexual dimorphism in the adult brain

As mentioned, the switch from female to male in zebrafish involves anatomical remodeling, a switch in sex-specific behaviors, and presumably remodeling of the corresponding brain regions (Dranow et al., 2016; Dranow et al., 2013). Microglia have been compelling candidate regulators of sex-reversal in the brain because they produce cytokines that promote neurodevelopment, including neurogenesis and oligodendrogenesis, and they have been proposed to shape neural circuitry but their role remains unclear (Casano and Peri, 2015; Lenz and Nelson, 2018; Li et al., 2017; Lopez-Atalaya et al., 2018; Nelson et al., 2019; O’Keeffe et al., 2025). Moreover, microglia are highly responsive to sex hormones, and the sex-specific differences that are established during development in mice are thought to affect brain function and microglia behavior in adulthood (Nelson et al., 2019). In zebrafish, estrogen receptors (ERs), particularly ERα and ERβ, are expressed by microglia, and estrogen signaling can influence microglial functions such as morphology, migration, and inflammation (Morale et al., 2006). In addition, some studies suggest that estrogen modulates microglial activity. For instance, estrogen signaling in microglia is thought to modulate inflammatory processes in the brain and neuroimmune responses after injury (Bruce-Keller et al., 2000; Villa et al., 2016). Androgen receptors (ARs) are also expressed by microglia in zebrafish (Gorelick et al., 2008), suggesting a potential role for androgens in neuroimmune responses, similar to the effects of estrogen. Sexual dimorphism in microglia numbers and regional colonization have been observed between the brains of male and female mice (Lenz and Nelson, 2018; Villa et al., 2018; Villa et al., 2019), and microglia ablation in young rats causes lifelong sex-specific behavioral deficits (Nelson and Lenz, 2017). Microglia also express prostaglandin receptors and produce prostaglandin (Minghetti et al., 1997; Minghetti and Levi, 1998) which when inhibited pharmacologically causes changes in microglia morphology and numbers (Lenz et al., 2013b; Nelson and Lenz, 2017). To determine if microglia distribution or morphology are sexually dimorphic in zebrafish, we used the microglia marker 4c4 to analyze microglia density in the Te, HT, and reticulospinal tract (RST) of adult brains. Although we observed no significant differences in the total number of cells in the brain areas analyzed in adult WT females and males (Fig. 5A and Suppl. Fig. 2A-2B), differences in microglia density were observed in some brain regions (Fig. 5B). Specifically, in adult female HT, 52% of the total cells were 4c4^+^ (microglia), while in males only 16% of HT cells were 4c4^+^ (Fig. 5B). Similarly, 31% of total cells in the RST of females were 4c4^+^ whereas only 18% were positive in males (Fig. 5B). This result indicates a higher density of 4c4^+^ cells in brain regions involved in sex-specific behaviors in adult females compared to males, which could account for the smaller size of these brain regions in females.

**Figure 5.**
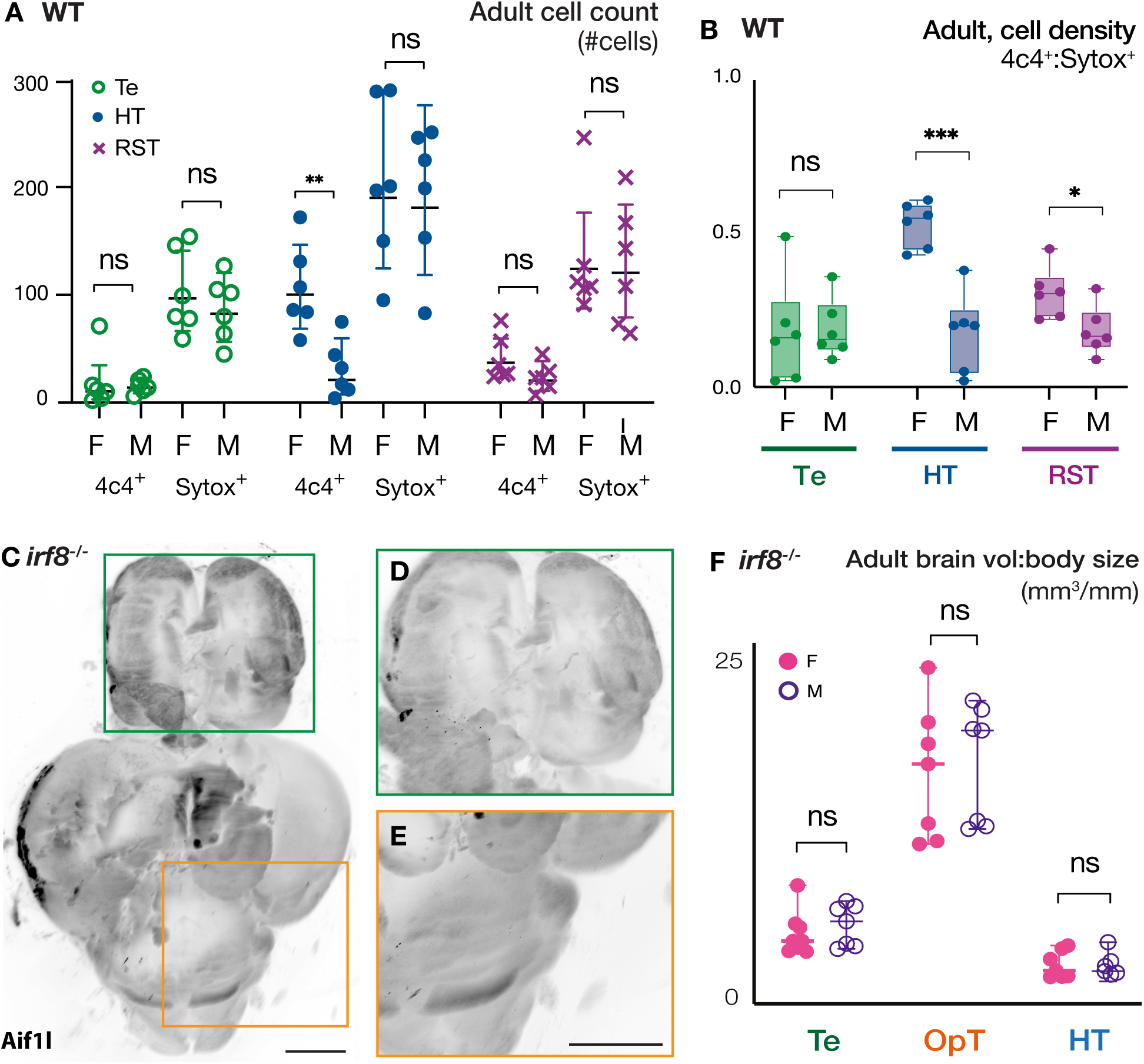
Sexual dimorphism of microglia density and regulation of region-specific brain volumes in adult zebrafish. **(A)** Number of Sytox (all nuclei) and 4c4 (microglia) positive cells in the indicated regions of adult WT brains. **(B)** Microglia densities in adult female and male brains. **(C-E)** Representative adult *irf8* mutant brain stained for 4c4 (microglia). Scale bars: 500um. Magnified boxed regions in **(D and E)**. **(F)** Volume/body quantification of indicated regions of adult *irf8^-/-^* brains devoid of microglia. Te: telencephalon; OpT: optic tectum; HT: hypothalamus; RST: reticulospinal tract. Statistical analysis for cell quantification: unpaired t-test with Welch’s correction and 95% CI. Statistical analysis for volume comparisons: ordinary Two-way ANOVA with 95% CI, and Fisher LSD test for multiple comparisons. P-values: ns≥ 0.05, *<0.05, ***<0.001.

Since microglia are known to be involved in neurogenesis, oligodendrogenesis, and cell proliferation (Casano and Peri, 2015; Lenz and Nelson, 2018; Li et al., 2017; Lopez-Atalaya et al., 2018; Nelson et al., 2019), they could contribute to shaping the observed sex-specific differences in the brain during development and/or adulthood. To determine if microglia contribute to sexual dimorphism of the Te, OpT, and HT, we analyzed the brains of females and males that lack microglia due to genetic ablation of the transcription factor *interferon regulatory factor 8* (*irf8*) (Ferrero et al., 2020; Kierdorf et al., 2013; Shiau et al., 2015b) (Fig. 5C-5E). In contrast to WT, no sexual dimorphism was observed between the Te, OpT, and HT of *irf8* mutant females and males (Fig. 5F). In addition to lack of sexual dimorphism in the brains of *irf8* mutants, we noticed that the ratio of brain volume to body size was greater in *irf8* mutants than in WT in the areas analyzed (Suppl. Fig. 2A-2C). This result suggests that Irf8-dependent cell lineages may play a broader role in maintaining the correct balance of cell types and numbers in the brain, potentially by contributing to neuronal death and clearance.

## Discussion

In sexually reproducing species, females and males develop distinct reproductive organs and physiological and behavioral features unique to their sex. Prior studies of adult zebrafish brains have identified differentially expressed genes between females and males as well as estrogen dependent differences in proliferation and cell death in the cerebellum and ventral telencephalon (Ampatzis and Dermon, 2007; Ampatzis et al., 2012; Sreenivasan et al., 2008). Similarly, prior studies identified dimorphically expressed genes and estrogen response elements (EREs) in whole brain microarray analysis (Santos et al., 2008). Comparison of gene expression profiles at juvenile and adult stages revealed few differences among juveniles but found modest androgen-associated differences in gene expression between adult male and female brains, leading to the model that sexual dimorphism of the brain is mediated by gonadal hormones (Arslan-Ergul et al., 2014; Lee et al., 2018b; Zhai et al., 2022). Among the genes with sexually dimorphic expression are regulators of reproductive and endocrine systems (Li et al., 2024; Santos et al., 2008; Sreenivasan et al., 2008). The detection of only modest differences in gene expression was postulated to be due to the plasticity of the female zebrafish brain to adopt male behaviors upon female to male sex reversal (Dai et al., 2023; Lee et al., 2018b). Despite modest dimorphism in gene expression, specific neuronal subtypes are sexually dimorphic in number and distribution between males and females (Lee et al., 2018b; Ogawa et al., 2021), and as in mammals, sex influences sensitivity and recovery from brain injury (Das et al., 2019).

Here, we investigated sex-specific morphology and plasticity of the brain in response to sex reversal. First, we found region-specific morphological differences between female and male adult brains that were observed only after sex determination and differentiation of the gonad. This result is consistent with the earlier reports that sexually dimorphic gene expression in the zebrafish brain emerges after sex determination and supports the hypothesis that gonad hormones sexualize the brain (Lee et al., 2018b). Finally, it defines the developmental window when sex-specific brain architecture is established. Specifically, with respect to brain areas activated during mating, including the telencephalon, optic tectum and hypothalamus (Li et al., 2024; Ogawa et al., 2021; Yabuki et al., 2016). In wild type, the optic and hypothalamic areas showed the greatest differences in size between female and male brains, consistent with previous studies showing cellular and gene expression differences in these areas between sexes (Gorelick et al., 2008; Ogawa et al., 2021; Waters and Simerly, 2009). Notably, sexually dimorphic expression of androgen receptors was seen in these brain areas of mice (Shah et al., 2004). That the same structures are dimorphic between sexes across species likely reflects the susceptibility of these regions to sexual hormones and their contributions to reproductive behaviors and function (Tsukahara and Morishita, 2020). Accordingly, sex hormone treatments, which have been shown to affect proliferation rates in adult brain regions, including the telencephalon and cerebellum, demonstrate their potential impact on overall size of these structures (Makantasi and Dermon, 2014).

We identified contributions of the tumor suppressor factor Chek2 to establishment of region-specific sexual dimorphism in brain volume. Specifically, we found that Chek2 is required for the smaller size of the female optic and hypothalamic areas. Given this finding and that loss of cells during development contributes to organization and function of many tissues, including the brain, it is likely that estrogen-dependent regulation of Chek2 maintains the smaller size of these brain regions in females (Ampatzis et al., 2012; Casano et al., 2016; Waters and Simerly, 2009; Xu et al., 2016b).

That Chek2 mutants are fertile, indicates that additional mechanisms contribute to sex-specific behaviors. Consistent with this notion, analysis of gene expression during sexual differentiation found that multiple pathways interact to regulate sexual behavior in teleosts, including upregulation of immune response factors during sexual differentiation (Lee et al., 2018a). Moreover, region-specific differences in microglia density were previously reported, with the highest in telencephalon, followed by the optic tectum, and finally the cerebellum of zebrafish adults; however, neither the sex of the fish examined nor whether microglia distribution or morphology patterns were sex specific was reported (Svahn et al., 2013). Consistent with the reported upregulation of immune response genes, we found sexually dimorphic distribution and densities of microglia between adult female and male zebrafish brains. Similar prominent sex-specific differences in microglia colonization, behavior and gene expression in response to estrogen or androgen exposure in development have been documented in mammals (McCarthy et al., 2015; Nelson et al., 2019). These sex-specific differences could be driven by differential colonization of microglia since, in vertebrates, immune cells are influenced by sex hormones and cell death (Casano et al., 2016; Loiola et al., 2019; Schwarz et al., 2012; Tränkner et al., 2019; Weinhard et al., 2018; Xu et al., 2016b). Moreover, hormone exposure in mouse developmental models indicates that the sex-specific neuroanatomy established during development persists and affects microglia function and behavior throughout life (Baker et al., 2004; Loiola et al., 2019; McCarthy et al., 2015; Nelson et al., 2019; Wu et al., 2016). In contrast, although microglia were dispensable for developmental establishment of sex-specific brain volume the larger brain volumes of mutants lacking microglia revealed microglia contributions to limiting the volume of the telencephalon and optic tectum of both sexes. Our findings and evidence from mammalian studies indicate that region-specific sexual dimorphism in microglia distribution and morphology is conserved, but that additional mechanisms, including cell death, contribute to regulation of sex-specific neuroanatomy of the adult brain in zebrafish.

Consistent with region-specific and/or redundant mechanisms contributing to sexually dimorphic neuroanatomy in zebrafish, Chek2 is required for sexual dimorphism in the telencephalon and hypothalamus but not the optic tectum. Considering that zebrafish undergo neurogenesis throughout their lifetime, microglia and Chek2-mediated pathways may differentially contribute to balancing proliferation/growth to maintain the smaller size of the adult female brain, similar to the situation during development in rodents (Byrd and Brunjes, 1998; Grandel et al., 2006; Oehlmann et al., 2004). Nonetheless, unlike mammals, reorganization of sex-specific neuroanatomy and behavioral pathways remains plastic in female zebrafish and can be modulated even in adulthood.

The observation that all *bmp15* mutant adult brains resembled non-mutant direct-differentiating male brains, suggests that the female brain retains plasticity and that female to male sex reversal involves activation of, or failure to repress, the mechanisms that promote male-specific neuroanatomy (Fig. 6). Because the gonad switches sex before secondary traits, it is likely that the shift from ovary to testes signals during gonad reversal and the resulting effects on sex hormone-responsive cell types promotes the region-specific sizes observed in direct-differentiating males. In mice, masculinization of the brain is blocked by feminizing factors (Nugent et al., 2015; Ahmed et al., 2008) Because ovary to testis sex reversal in *bmp15* mutants occurs relatively early in reproductive life and mutant oocytes fail to reach mature stages, the concentration and/or duration of ovary signals may not be sufficient to counteract masculinization (Bravo et al., 2023; Dranow et al., 2016; Zhai et al., 2023). Consistent with this notion, Bmp15 deficiency has an indirect effect on estrogen production and estrogens are known to modulate region-specific brain growth during development in mammals (Ahmed et al., 2008; Nugent et al., 2015; Zhai et al., 2023). Although juvenile brains were not compared, given that adult mutant brains were larger than those of male and female siblings, it is possible that juvenile mutant brains directly acquired male-like characteristics in response to the estrogen deficits and consequent rise in signals that drive masculinization.

**Figure 6.**
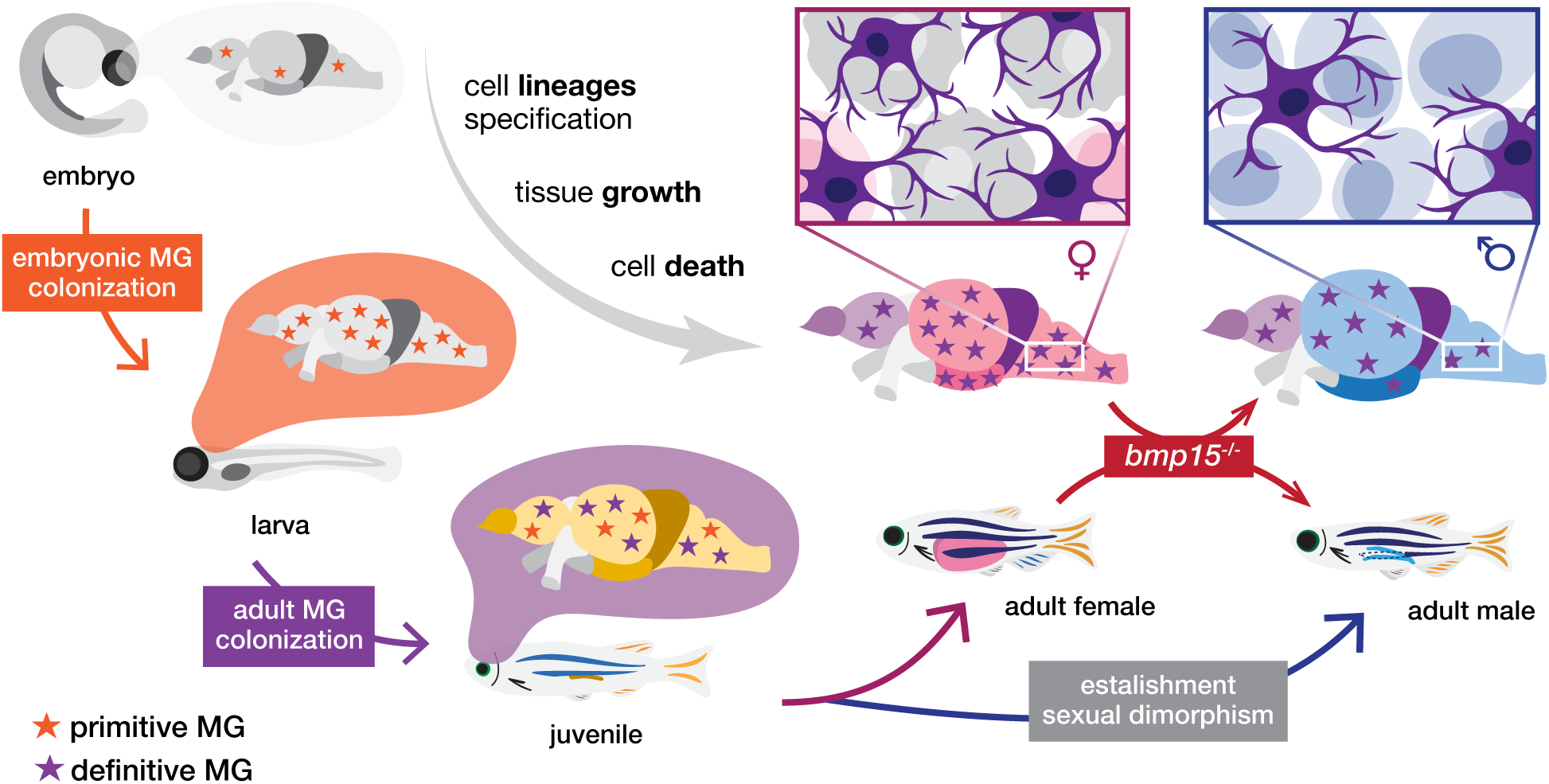
Microglia and cell-death contributions to sexual dimorphism in the adult zebrafish brain. Schematic illustrates the microglia colonization for the embryonic (primitive), and adult (definitive) waves of tissue resident macrophages. During initial development, the brain grows, and distinct cell lineages are specified. During the late juvenile stage (around 28dpf), sex-specific traits arise between female and male brains. Differences in microglia density (higher in females than males), and in brain structure volumes (larger Te, OpT, and in males).

Zebrafish adult females retain the plasticity to switch primary and secondary sex from female to male as evidenced by sex-reversal of *nanos3* mutant females. However, the adult female brain may become less plastic with age as we observed no difference in brain region size between sex reversed *nanos3* mutants and heterozygous sibling females. Nonetheless, sex reversed *nanos3* mutants acquire male mating behaviors, suggesting that in adulthood these behavioral pathways are remodeled through a mechanism that doesn’t involve overt anatomical changes of the brain, but could involve modulation of microglia numbers or activity or modulation of cell type composition within the brain.

Although *irf8* mutants could develop as either sex, both sexes had larger brains compared to siblings and no sexual dimorphism was observed between sexes and mutant males and females were fertile in natural mating trials. Thus, microglia do not seem to be required for reproductive function in direct differentiating males and females. However, it is possible that microglia are involved in remodeling and acquisition of male-specific behaviors after sex reversal. Unfortunately, the available genetic models eliminate or block the activity of all circulating and tissue resident macrophages and completely block sex-reversal of the gonad and secondary sex traits (Bravo et al., 2023). Thus, specific contributions of tissue-resident macrophages in sex reversal, including microglia, remains challenging to resolve. Addressing this question awaits development of methods to ablate microglia without disrupting circulating or other tissue resident macrophage populations. Interestingly, although Chek2-dependent pathways are required to establish sexually dimorphic neuroanatomy and sexual dimorphism in regional microglia density, neither Chek2 nor Irf8 are required for direct sex differentiation or reproductive behaviors, since fish lacking Chek2 or microglia develop and behave normally as females or males. This observation suggests that acquisition of sex-specific behavioral pathways (e.g. mating) during development is either not regulated by Chek2-dependent pathways or microglia, or is redundantly regulated by Chek2, microglia, or other factors. Although transcriptomic approaches have been performed to compare adult female and male brain areas, a gap remains in our understanding of molecular differences at the level of individual cells within the mature brain (Chen et al., 2019; Sagi et al., 2024). Filling this gap should enhance our understanding of the basis of sex-specific susceptibilities within the nervous system and has potential to reveal biological and therapeutic targets.

In mammals, development, maintenance, and remodeling of reproductive tissues are strongly influenced by peaks and changes in sex hormones that occur in all sexes at various timepoints throughout life (Ahmed et al., 2008; Gegenhuber and Tollkuhn, 2020; Mosconi et al., 2024). These shifts of hormones influence the function of many tissues in the body, with susceptibility of the nervous and immune systems being the most appreciated (Bereshchenko, 2018; Klein et al., 2016; McCarthy et al., 2015; Sciarra et al., 2023). Many neuropsychiatric disorders that are sex biased arise or are diagnosed near puberty, pregnancy, or menopause –all periods of major hormonal change in persons with ovaries (Craig and Murphy, 2007; He et al., 2024; Mathews et al., 2015). The susceptibility of the brain to hormonal changes (Ahmed et al., 2008; Geleta et al., 2024; McEwen and Milner, 2017; Pradhan and Olsson, 2015), or to acute loss of gonad signals, could influence the severity or onset of sex biased neurological diseases. Whether loss of Chek2 or microglia dysfunction influences other physiological processes in the brain, including responses to injury, inflammation, or aging has important implications for normal and pathological physiology states of sex hormone responsive cells, sex hormone-driven tissue remodeling, and sex biased disorders.

### Data Availability

All data and analyses are contained within the manuscript. There were no sequencing or related datasets generated in this study.

## Supporting information

Supplemental Figures for Bravo and Marlow

## Acknowledgments

The authors thank former and current members of the Marlow Lab for valuable discussions, the Center for Comparative Medicine Staff for fish care, and the Microscopy and Advanced Bioimaging CoRE at the Icahn School of Medicine at Mount Sinai for microscopy assistance. This work was supported by startup funds to F.L.M. and National Institute of General Medical Sciences (NIH-R01-GM089979 and NIH-R35 GM153244 to F.L.M.).

## Author contributions

Conceptualization, P.B. and F.L.M; formal analysis, P.B. with guidance from F.L.M.; investigation, P.B.; visualization, P.B.; supervision, F.L.M.; project administration, F.L.M.; funding acquisition, F.L.M.; writing, critical review, and editing of the manuscript, P.B. and F.L.M.

