## Supplemental Figures for Bravo and Marlow for "Microglia and Chek2 contribute to sex-specific organization of the adult zebrafish brain"

### Supplemental Figures and Legends

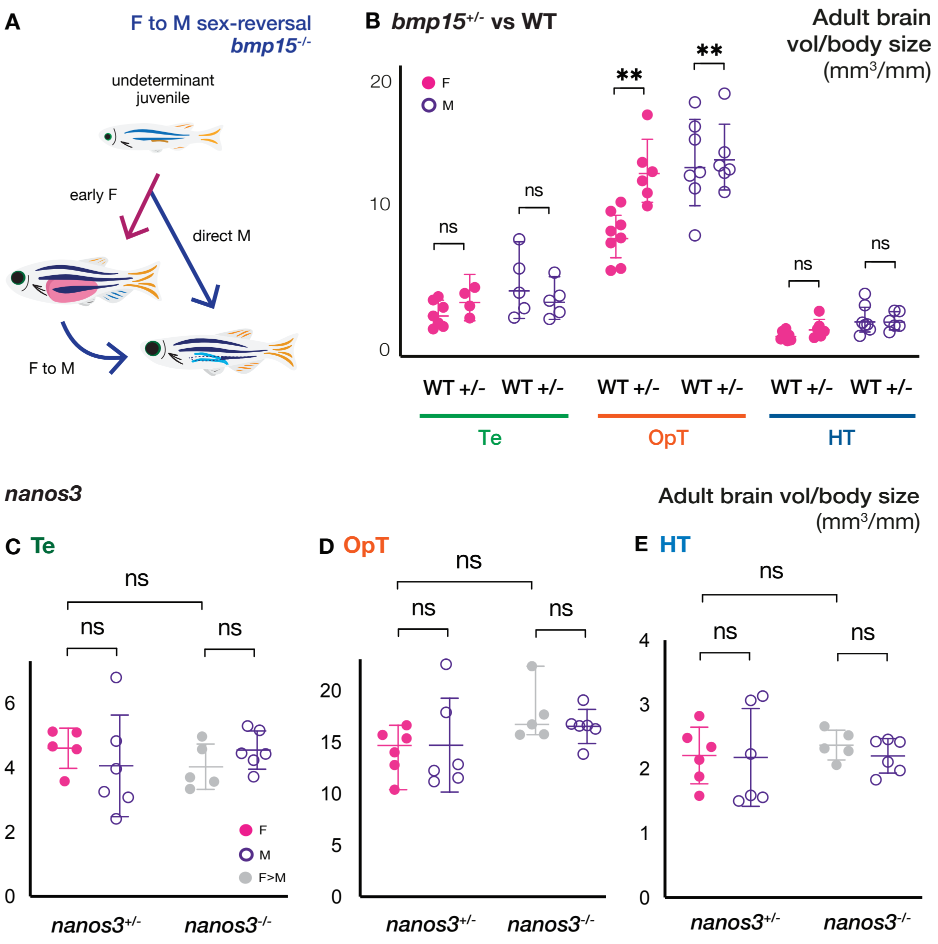


**Suppl. Figure 1 Anatomical comparison of brains of sex-reversal mutants to WT**

**(A)** Schematic depicts indeterminant juvenile stage and subsequent sex reversal associated with loss of *bmp15* . **(B)** Comparison of indicated brain region volumes between WT and *bmp15*$^{+/-}$ adult females and males. **(C-E)** Comparison of **(C)** Te, **(D)** OpT, and **(E)** HT brain volumes to body size of adult *nanos3* heterozygous and homozygous mutants. Te: telencephalon; OpT: optic tectum; HT: hypothalamus. Statistical analysis: ordinary TWO-way ANOVA with 95% CI, and Fisher LSD test for multiple comparisons. P-values: ns≥ 0.05, **<0.01.

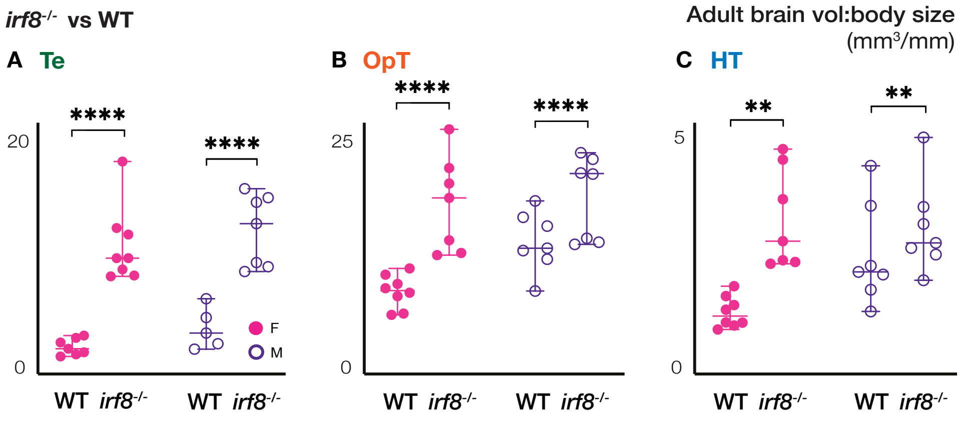


**Suppl. Figure 2 Anatomical comparison of *irf8* mutant and WT brains**

**(A-C)** Volume/body size comparison of *irf8*$^{-/-}$ and WT female and male brains **(A)** Te, **(B)** OpT, and **(C)** HT. Te: telencephalon; OpT: optic tectum; HT: hypothalamus. Statistical analysis for volume comparisons: ordinary Two-way ANOVA with 95% CI, and Fisher LSD test for multiple comparisons. P-values: **<0.01, ****<0.0001.
